# SecSel, a new software tool for conservation prioritization that is applicable to ordinal-scale data for multiple biodiversity features

**DOI:** 10.1101/2021.02.15.431247

**Authors:** Takenaka Akio, Oguma Hiroyuki, Amagai Yukihiro, Ishihama Fumiko

**Affiliations:** National Institute for Environmental Studies, Onogawa, Tsukuba, Ibaraki, Japan

## Abstract

SecSel, a protected-area prioritization tool, has been developed to help design areas that efficiently protect multiple features, including conservation of biodiversity and use of ecosystem services. The prioritization by SecSel is based on evaluation of the local units of each feature. The evaluation metrics should be quantitative but need not be ratio scale. The minimum requirement of the input data is that they are ordinal. The conservation target is the number of local units with high values of each feature to be protected in the area. SecSel can handle conflicts among features, including conflicts between conservation and utilization of land or specific ecosystem functions. Before the selection procedure, one of a conflicting pair of features in a site is discarded. That decision is based on the dispensability of the local unit to fulfilling the conservation target of each feature. SecSel also considers the cost of including each site in the protected area and the compactness of the area in terms of total boundary length or the distance to the nearest site. To demonstrate the functionality of Secsel, we used it to design land use in an alpine region of northern Japan where conservation of alpine vegetation and its recreational use are important considerations.

## Introduction

Prioritization of conservation areas is one of the major conundrums in conservation ecology [1-3]. Various methods have been proposed for the prioritization of local conservation units and the design of regional conservation areas [3-6].

Appropriate design tools are chosen on the basis of available data and the purposes of the prioritization analysis. When detailed data are available, the protected area can be designed based on a quantitative evaluation of conservation effects. With less informative data, the evaluation of conservation effects is inevitably less quantitative. Binary presence-absence data for organisms, for example, do not include the quality of local populations. Inclusion of any local population is considered to contribute equally to the conservation of target species.

Both of the widely used conservation prioritization tools, Marxan [5] and Zonation [4], require ratio-type evaluation data for biodiversity features, which can be summed and subtracted. Marxan set conservation target as the sum of the evaluation data for each feature to be protected [5]. If the target exceeds the summed value, the difference is considered as a cost component. In Zonation, the prioritization process starts from the largest protected area covering a whole region [4]. The least valuable site is initially omitted from the protected area. For a given biodiversity feature, the value of a site is the fraction of the total amount of the feature included in the protected area. The functionality of these algorithms requires that the evaluation data be ratio-scale for both Marxan and Zonation.

Quantitative data such as area covered by vegetation to conserve, population size, population density, and biomass meet this requirement. Binary data such as presence-absence, which provide limited information, also can be handled as ratio-type data and fed to Marxan and Zonation.

It is often the case that non-ratio type data that provide more information than presence-absence data are available. A typical example is a rough evaluation of population size or quality (e.g., Braun-Blanquet survey methods cover classes of plants in a phytosociological survey [7]). For example, experts are able to distinguish between stable populations with high density, populations with lower densities, and accidental observations of a few individuals. Such data cannot be added or subtracted, but they are more informative than presence-absence data.

SecSel is a newly developed prioritization tool designed to handle rank-type data for prioritization analyses. SecSel selects areas for conservation that include local units that are highly ranked on the basis of multiple biodiversity features. Ecosystem functions and other purposes of land use may also be included in the analysis. In SecSel, a set of several local units with low ranks for a biodiversity feature is not a substitute for a single unit with a high rank. This limitation causes some inflexibility in site selection, but flexibility remains as long as there are ties between local units. Such ties occur in many cases when biodiversity features are roughly estimated.

In many cases of conservation planning, conflicts between protection and other land use are inevitable. There are several approaches to finding compromises when such conflicts arise. For example, Chan et al. (2011)[8] have compared two approaches to incorporating ecosystem services in a Marxan analysis: handling the services as benefits to be targeted by conservation and as co-benefits or opportunity costs. With Marxan with Zones [6], which is an extended version of Marxan that can treat multiple zones for different objectives, land management uses, including biodiversity conservation, are categorized into different zones. With another widely used prioritization tool, Zonation, conflict may be simply resolved by masking out an area for land use other than biodiversity conservation. In the advanced version of Zonation, areas suitable for land uses that have negative impacts on biodiversity are given negative weight in the ranking of priorities so that these areas are eliminated from the ones to be protected in the early stage of site selection[9].

In SecSel, ecosystem functions are also features to be considered. It is sometimes the case that use of land to exploit ecosystem functions conflicts with conservation of some of the biodiversity features. In addition, other conflicts may occur between biodiversity features. For example, there may be pairs of biodiversity features for which conservation of one has a negative impact on the other. Examples include species of woodland animals and grassland animals, or aquatic species in eutrophic habitats and those in oligotrophic habitats.

There are several approaches to resolving such conflicts. One approach is allocating an area for commercial land use beforehand and subsequently selecting an area for conservation from the remaining land. Such an approach is simple, but it may lead to deterioration of areas that are irreplaceable for the conservation of some features of biodiversity. Chan et al. (2011)[8] have used a widely used prioritization tool, Marxan, to try to incorporate the use of ecosystem services that may be positively or negatively correlated with conservation of biodiversity into site selection. In their analyses, ecosystem services are handled as targets to be conserved, or they are regarded as co-benefits or costs associated with the conservation of biodiversity features.

Before site selection, SecSel searches for conflicts between local units with incompatible features within sites. For every conflict, which of the incompatible local units has higher priority is determined. Features with lower priority are ignored in the subsequent selection procedure. The priority is greater when the local unit in the site has a relatively high rank. Details are described in section *Conflict*.

This paper provides an introduction to the basic purpose and rationale for SecSel. As an example of its application, we applied SecSel to the design of land use in an alpine region of northern Japan where consideration was given to conservation of several biodiversity features and their recreational use.

## Methods

### Software description

SecSel is coded in Python 3. It is open source and is available from the website (http://www.nies.go.jp/biology/en/data/tool/secsel/index.html). Its function is controlled by a setting file, which is a text file that describes data file names and parameter settings. In the next sections, we define the basic terms used by SecSel, explain the required input data, outline the selection process, and describe the output data. A graphical summary of these workflow is provided in Fig. 1.

**Fig. 1.**
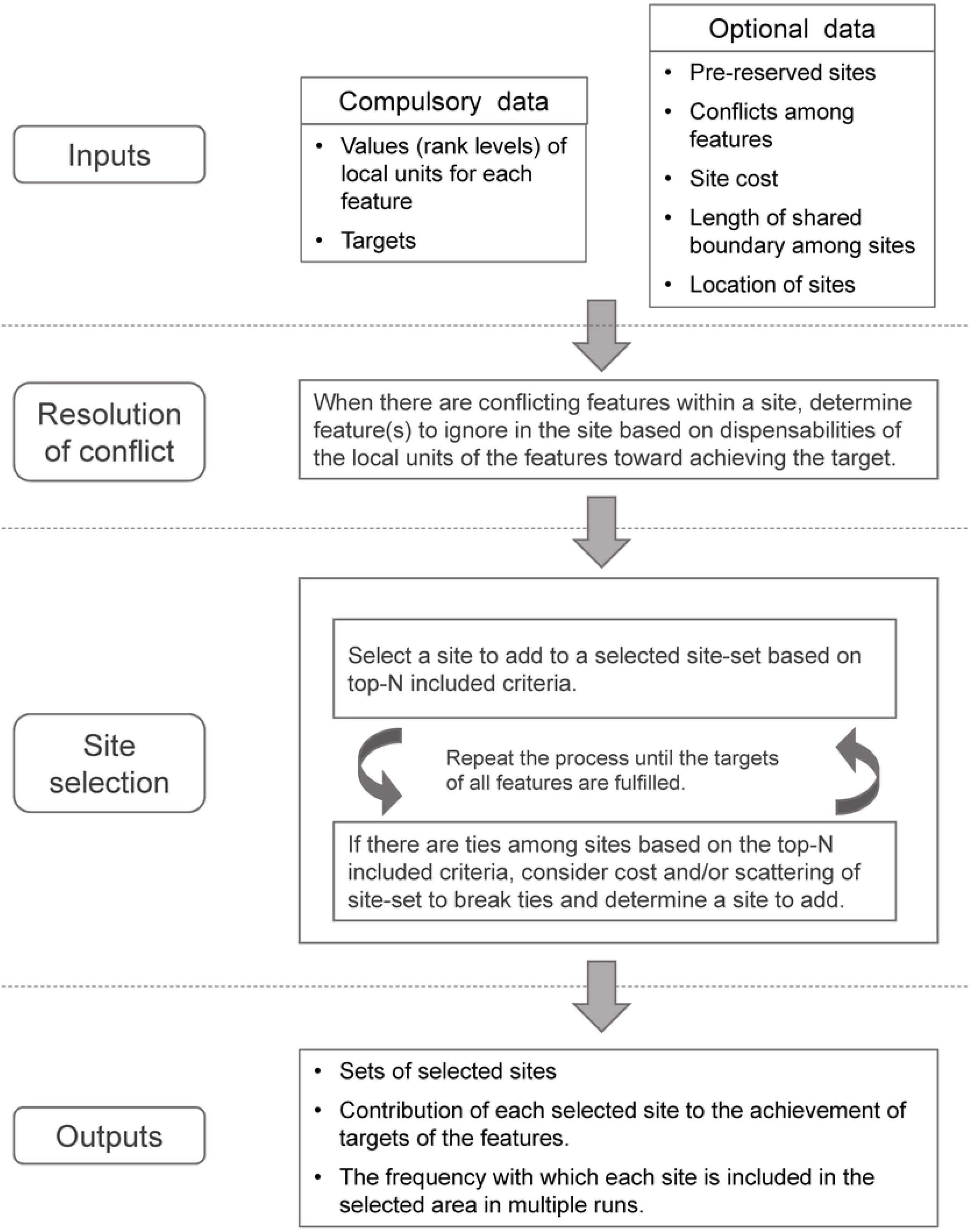
A graphical summary of spatial prioritization process by SecSel.

### Definitions of terms used in SecSel

Abbreviated explanations of basic terms used in SecSel are provided in Table 1. Detailed definitions and explanations are in the following sections.

**Table 1.**
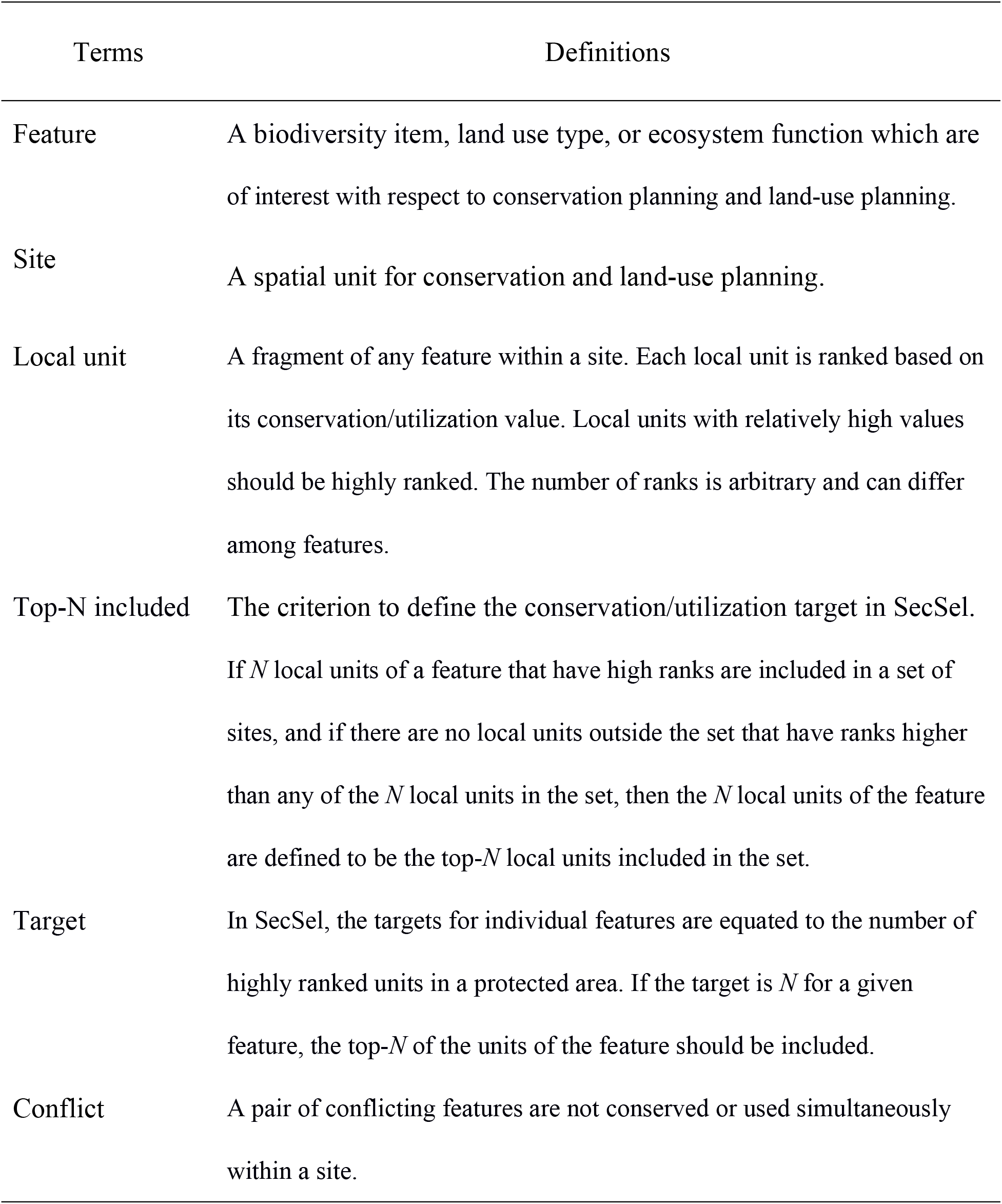
Abbreviated explanations for the definitions of terms used in SecSel. See the main text for details.

### Feature

A feature is a biodiversity item, land use type, or ecosystem function at any level. Biodiversity features include taxonomic classification at any level as well as the types of vegetation, habitat, and ecosystem. Land use types and ecosystem functions include functions at any level, including agricultural and silvicultural land use. Any land use not related to ecosystems, such as urban areas and power plants, may also be included in the analysis.

### Site

A site is part of an area within a region targeted for conservation and land-use planning. Sites are not necessarily of the same size, and they do not need to be spatially continuous. Scattered ponds or fragments of woodlands or grasslands are example of separated sites. Sites in SecSel correspond to the planning units of Zonation [4].

### Local unit

A local unit is a fragment of any feature within a site. In other words, each site includes local units of features of interest with respect to conservation planning and land-use planning. Each local unit is assigned to an arbitrary rank level for the analysis by SecSel. Local units with relatively high conservation values should be highly ranked to conserve biodiversity features. Units that are better suited for a use or that provide more services should be ranked highly to facilitate use of land and ecosystem functions. The number of ranks is arbitrary and can differ among features.

### Top-N included

If *N* local units of a feature that have high ranks are included in a set of sites, and if there are no local units outside the set that have ranks higher than any of the *N* local units in the set, then the *N* local units of the feature are defined to be the top-*N* local units included in the set. For example, if three local units of a feature within a set have been assigned ranks of 4 or 3 (4 is the highest) and the ranks of other local units are 3, 2 or 1, then the set is said to be top-3 included. This criterion is introduced to define the conservation/utilization target as described in the next section.

### Target

In SecSel, the targets for individual features are equated to the number of highly ranked units in a protected area. If the target is *N* for a given feature, the top-*N* of the units of the feature should be included. Based on this concept, the local units of the feature should be top-*N* included in the protected area. If the ranks are highly resolved and no local units have the same rank, the set with the minimum number of sites that fulfill the top-*N* included criterion of a feature is unique. If the resolution of the ranks is relatively low and ties are frequent, the selection of sites will be more flexible. This flexibility makes it possible to identify a compact set of sites that fulfills all the targets for multiple features.

### Conflict

In SecSel, a conflict may occur in a pair of local units of features within a site. Local units of conflicting features are not conserved or used simultaneously within a site. Conflicts between incompatible features within sites are resolved before site selection. The local unit that is more critical for achieving the target of the feature has higher priority in resolving the conflict. A local unit of a feature is given lower priority than more critical units of other features. Because an explicit decision is made about which feature is given priority at a site, it is clear which features within a selected site are to be protected/used in the implementation of conservation design. The details of the decision process are described in section **Resolution of conflicts**.

### Input data

#### Compulsory input data

For each feature to be considered, data of evaluation for individual local units are necessary. Furthermore, there should be targets for individual features.

#### Rank levels of local units

Evaluation of local units within each site should be prepared for each feature. For the features to be protected, the evaluation is based on the conservation value. For the features to be used (e.g., ecosystem functions and various land uses), the evaluation is based on the quality and/or quantity of the benefit the site would provide.

The evaluation is quantified on an ordinal, interval or ratio scale. Binary data consist of 1 for presence and 0 for absence; three levels may be high, low, and no value; or there may be continuous values that describe the area of concern in an ecosystem or the size of a population. All these metrics meet the requirement of SecSel. There is no limitation on the resolution of the evaluation. On the one hand, the higher the resolution, the more precise the selection of the highest priority local units will be obtained. On the other hand, low resolution enables more flexible selection of local units. Flexibility makes it possible to select a compact set of sites to achieve targets for multiple features. Flexibility also facilitates selection of multiple protected areas that achieve almost the same conservation efficiencies.

#### Targets

For every feature, the number of top local units to be included among the selected sites is specified as a target. For biodiversity features, the target is a conservation requirement. For features to be used, the targets are utilization requirements. If the target of feature *i* is *N*_i_, the target is achieved when the local units of this feature in the selected sites satisfy the top-*N*_i_ included criterion. It should be noted that collection of local units of low value is not a substitute for a highly prioritized local unit. Targets may or may not be the same among features. A target can be assigned to each feature independently.

#### Optional input data

In addition to the above-mentioned compulsory input data, SecSel can consider pre-reserved sites, conflicts among features, cost of conservation, and compactness of conservation area in the site-selection process.

#### Pre-reserved sites

Sites that should be included in the protected area can be specified as pre-reserved sites. When the current protected area is to be expanded, sites included in the present protected area are specified as pre-reserved sites. SecSel decides which sites should be added to these pre-reserved sites.

#### Conflicts among features

As described above, conflicting features are those that cannot be conserved/used simultaneously in a site. To consider conflicts and avoid incompatibilities, data about conflicting pairs of features should be provided. SecSel decides which feature has higher priority and which has lower priority at each site where there is a conflict among local units of two or more features. Section **Resolution of conflicts** describes the algorithm of the decision.

#### Site cost data

Secsel can consider the cost of including individual sites in the protected area. For example, the cost can be the amount of money paid to obtain the land and/or to perform conservation management in the site. Given cost data for individual sites, SecSel tries to find a set of sites with the lower cost. Like the data of the conservation/utilization values of local units of features, the cost data are on an ordinal scale with no specified degree of resolution. The cost data are used to determine which of the candidate sites of similar conservation efficiency should be selected. Subsection *Cost considerations* explains the details of the cost consideration process.

The cost of a site may depend on which feature is to be conserved/used in the site. SecSel does not handle such multiple cost criteria. If a special cost should be paid for a feature, the evaluation of the local units of the feature may be decided on the basis of the cost of conservation/utilization of the feature within each site.

#### Shared boundary data

The length of the boundary shared by two sites can be used to select the set of sites that is less scattered. Like cost data, the length of the boundary is used to determine which of the candidate sites of similar conservation efficiency should be selected. There is no guarantee that the selected set of sites has the smallest perimeter.

### Outline of site-selection processes

#### Basic process of site selection

In the first step of the selection process, sites are selected to satisfy the top-1 included criterion for all features. Once that criterion is satisfied, sites are added to fulfill the top-2 included criterion for all features for which the target is 2 or larger. This process is continued by incrementing the number of top local units to be included. The process is terminated when the targets are fulfilled for all features of interest.

For each step of selecting a site to add, a greedy algorithm is employed. For each unselected site, the number of features that would fulfill the top-*N* included criterion (*N* being incremented during the selection procedure) is calculated. The site with the maximum number of features that newly fulfill the top-*N* included criterion is the site to be added to the sites that have already been selected. If there are ties, one site is selected randomly.

#### Cost considerations

When cost is to be considered, it is used to break ties between sites in the number of features that newly fulfill the top-*N* included criterion. To place more emphasis on reducing cost in the selection process, candidate sites may include sites that are not necessarily among the sites that most effectively increase the number of features that fulfill the top-*N* included criteria. The number of candidate sites, including the less efficient but less costly sites, reflects the emphasis on cost reduction. The larger the number of candidate sites, the greater the odds that a less efficient site may be selected because of its small cost. This process may increase the number of sites required to achieve the targets of all features. The final set of sites is not necessarily the set of sites that minimizes the total cost irrespective of the number of candidate sites. This result is an inevitable consequence of a heuristic algorithm.

#### Selecting less-scattered sites

A less-scattered set of sites can be selected in SecSel. SecSel implements two methods for this purpose.

One of the methods considers boundary length. During the process of site selection, the total boundary length of the set of selected sites changes. It typically, but not necessarily, increases. The change in length by addition of a site depends on the geometrical relationship of the additional and already-selected sites. Addition of a site that shared a boundary with already-selected sites would increase the total boundary length less than addition of an isolated site. Thus, by considering the change of the total boundary length, sites adjacent to already selected sites are more likely to be selected. To break ties in the selection of sites based on the number of features newly fulfilled in the top-*N* included criterion, the site that causes the smallest increment to the total boundary length is selected.

The other method considers the distance of a site to the nearest site among the other selected sites In this method, the distance to the nearest selected site is used to break ties in the same way that the boundary length method is used in the site-selection process. The site that lies the shortest distance from the selected site is chosen. The merit of this method is that it is applicable in a case when the sites naturally do not share a common boundary (e.g., each site is a different pond or a fragmented patch of forest). This method helps to avoid selecting isolated sites that are distant from already-selected sites, and it is expected to reduce the maximum of “the distance to the nearest sites” within the conservation area.

#### Consideration of both cost and scattering

In the consideration of both the cost and scattering of sites, scattering is the first criterion used to break ties at each step of site selection. If there are still ties after consideration of scattering, then cost is considered. In the early stages of the site-selection processes, sites that are highly efficient with respect to fulfilling the top-*N* inclusion requirement are not likely to be adjacent to each other as long as there is spatial autocorrelation in the distribution of each feature. Neighboring sites are likely to have more similar compositions than randomly selected sites, and they are less likely to complement each other. Scattering is therefore not likely to determine which of the tied sites should be selected. If there are differences in costs among sites, those differences could decide which site is selected. As more and more sites are selected, it becomes increasingly likely that candidate sites will share a boundary with or close to already-selected sites. The protected area thus expands by adding adjacent sites to low-cost, previously selected sites. If there is spatial autocorrelation of costs, the costs of the sites added to the low-cost sites are also expected to be low. The protected area is therefore likely to start from one or a few core areas in a low-cost region; adjacent sites are then added to expand the core areas. The final result is a less-scattered set of selected sites in low-cost regions.

Consideration of costs and scattering may lead to a protected area with a relatively large number of sites after the heuristic selection process. The number of sites varies between cases. If there are relatively large numbers of highly ranked local units with numerous features, there is more flexibility in site selection to achieve targets, and consideration of costs and scattering would have less of an effect on the final size of the protected area.

### Resolution of conflicts

When there are conflicting features within a site, Secsel determines which one of the conflicting features is to be ignored prior to site selection. To determine which feature is to be ignored, the dispensabilities of the local units of the individual feature toward achieving the target are compared. To enable this comparison, a dispensability index, *S*, is defined as follows:

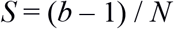

where *N* is the target of the feature, and *b* is the position of the local unit within all local units of a feature in the descending order of value. If there is more than one local unit with the same rank, *b* is calculated as follows:

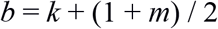

where *k* is the number of local units of the feature with higher ranks, and *m* is the number of local units with the same rank as the unit in the focal site. For example, if the top four local units have the same rank, all units are assigned *b* equal to the mean of 1–4, i.e., 2.5. If there are five rather than four tied local units, *b* is 3.0. A larger *b* results in a larger dispensability index, *S*. When more local units are available as substitutes for the one in the focal site, the local unit becomes more dispensable.

The value of *S* varies inversely with the target, *N*. The implication of a larger *N* is that the local unit in the focal site is less dispensable. Even when the *b* is the same among features, the one with the larger *N* requires more local units with high ranks, and the local unit in the focal site is thus less dispensable.

After the *S* values of conflicting local units within a focal site have been calculated, the local unit with the larger *S* is ignored. Because the omission of a local unit affects the *S* values of other units of the feature, the *S* values of remaining units are recalculated. The fewer local units that remain, the less dispensable will be the remaining local units.

#### Outputs

The output of SecSel is a set of selected sites listed in the order of selection. Additional information includes the features by which each site contributed to the achievement of targets. When conflicts among features are taken into account, the information used to ignore some features is provided. This information is crucial to deciding conservation/utilization practices within selected sites when there are local units of conflicting features.

In addition, SecSel outputs the frequency with which each site is included in the selected area in multiple runs. There are some stochastic processes in the SecSel site selection. First, ties can occur when deciding which site contributes most to achieving targets. One of the sites that is tied is chosen randomly. Such ties occur frequently, especially when there is a low resolution of the local unit evaluation. It should be noted that low resolution leads to flexibility with respect to site selection, and it widens the range of variation of site selection in the process of tie breaking. There is another stochastic process in the resolution of conflict. When there is a tie in the dispensability index among local units with conflicting features, one of the units is randomly given priority. Ties can also occur in site selection during consideration of cost and scattering.

A result of these stochastic processes is that site selection can vary between replicated runs of SecSel. The range of variation depends on the nature of the given data. The greater the probability of ties, the more likely that the variation of selected sites is large.

SecSel automatically repeats the user-specified number of runs. The frequency with which each site is included in the selected area in multiple runs may be an indicator of the irreplaceability of the site. However, a combination of sites that are frequently selected does not necessarily result in protected areas that efficiently fulfill targets. This is because the sites in the set chosen in a single run complement each other to achieve the goal, and there is no guarantee that the goal will be achieved if the set is broken apart. Such properties are common to complementarity-based prioritization, and selection frequency results should be interpreted with caution in this regard.

## Example of site prioritization

### Case description: An alpine vegetation in northern Japan threatened by climate change

The Taisetsu Mountains in northern Japan are the site of the largest national park in the country (Fig. 2). Many alpine plant species (365 species) grow in the Taisetsu Mountains, including subspecies and variants as well as 27 Japanese endemic species [10-12]). In addition, alpine vegetations are expected to be particularly vulnerable to climatic changes [13]. Thus, alpine vegetation should be highly prioritized for conservation.

**Fig. 2.**
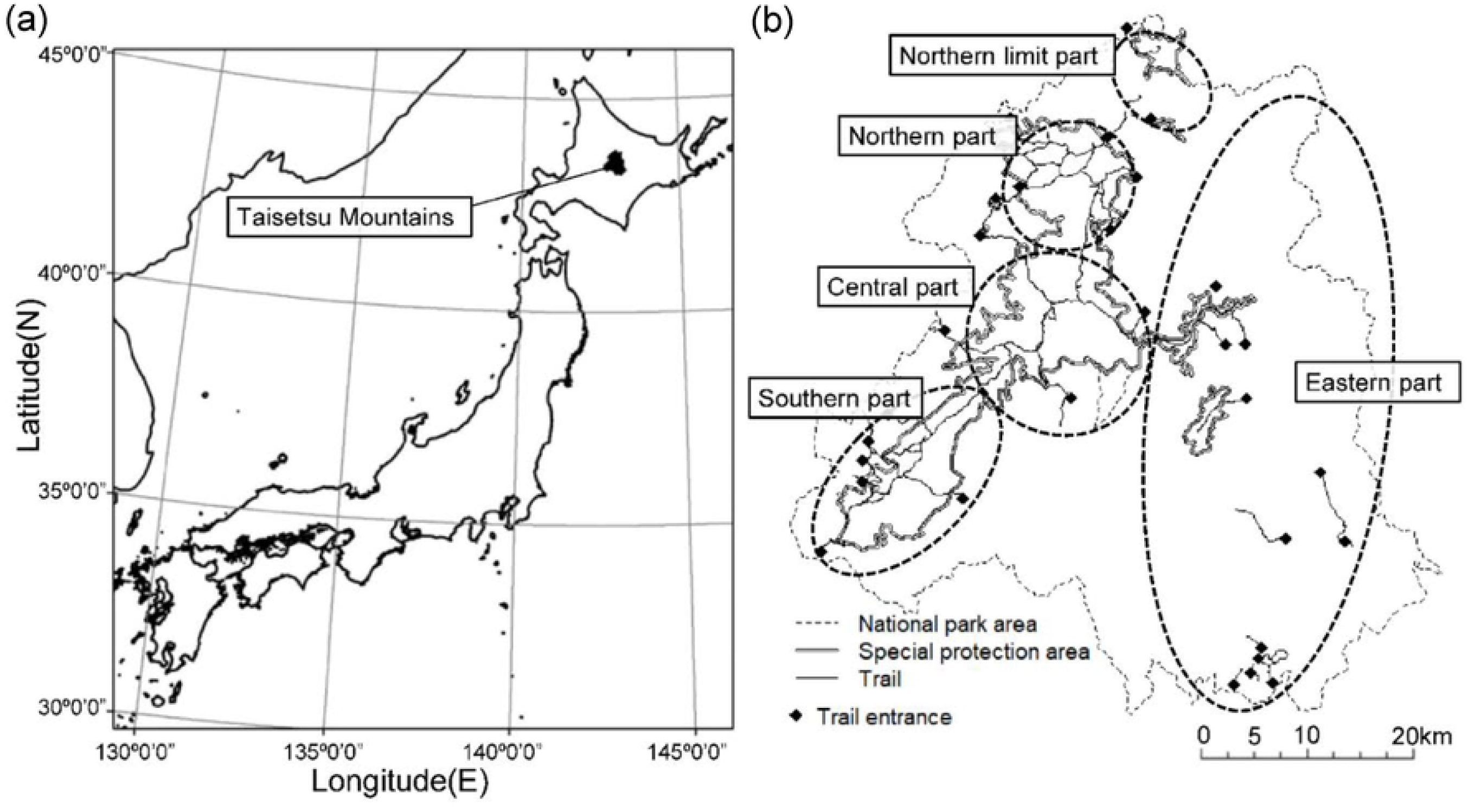
Study site. (a) The position of Taisetsu Mountains and (b) the national park area.

National parks in Japan are expected to play roles not only in biodiversity conservation but also in the provision of ecosystem services. Fields of alpine flowers attract visitors and are one of the most important resources for tourism.

### Evaluation of biodiversity features

We evaluated the value for conservation based on the area of alpine vegetations which are the targets of biodiversity conservation. We classified alpine vegetations into four types—snow meadow, fellfield, wilderness, and shrubs—for purposes of considering conservation of biodiversity. Geographic Information System data at the 1:25,000-scale were used [14] to calculate the area of each alpine vegetation type in each grid cell. The grid cells were approximately 1 × 1 km (30-second latitude difference and 45-second longitude difference). We then categorized the habitat quality of each grid cell for each vegetation type into four rank classes based on the area of the vegetation type within the cell (Fig. 3).

**Fig. 3.**
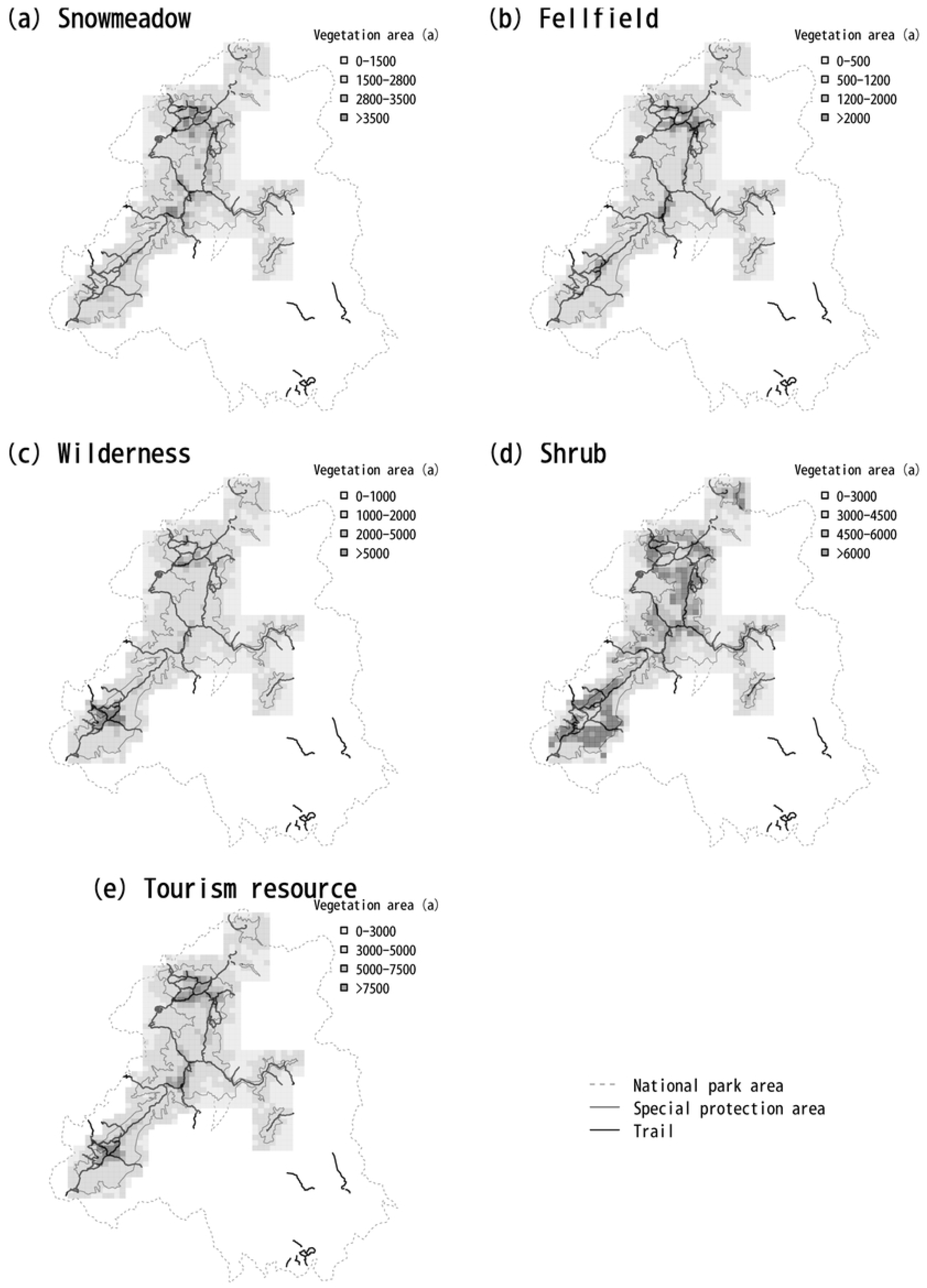
Distribution of each type of vegetation in Taisetsu National Park.

### Evaluation of ecosystem services

We measured the value of ecosystem services from the viewpoint of tourism resources. We considered the sum of the areas of snow-meadow and fellfield vegetation as the quality of tourism resources, because these types of vegetation form attractive flower fields, which are highly valuable resources for tourism. We categorized the resource quality of each grid cell into four rank classes based on the sum of the areas of the vegetation types within the cell (Fig. 3). These vegetation types were also included in the targets of biodiversity conservation, but we considered conservation and tourism to be competitive uses, and we selected sites for tourism and sites for conservation separately.

### Conflicts among features

Tourism can be a threat to the conservation of vegetation because of trampling and illegal digging. We therefore considered conservation and tourism to be competing uses of a site. Among conservation features, alpine shrubs out-compete other types of alpine vegetation, and we considered that there were conflicts between these vegetation types.

### Costs of conservation management

Options that we considered for conservation management included patrolling to prevent illegal digging, monitoring changes in the states of vegetation, and cutting back sasa-bamboo and shrubs, which would out-compete alpine vegetation in a climate change scenario. Accessibility was an important consideration for all forms of conservation management as well as for tourism. We thus considered the time required to reach each cell as a cost in the analysis. To calculate the required time, we first calculated the distance from the trail entrance to a cell along the climbing trail. If a cell did not include a trail within it, we also calculated the distance from the nearest trail to the cell. We then calculated the time required by assuming that it took 54 min to walk one kilometer along the trail and 120 min outside the trail.

### Conservation targets

We set a conservation target for each biodiversity feature and ecosystem service in eight cells. We performed 100 iterative calculations, and we summarized the frequency of selection for each cell.

## Results

Without consideration of cost or scattering penalty (cost-off and boundary-off in SecSel), sites in the northern and southern parts of the Taisetsu Mountains were often selected as a whole, and sites in the central part were selected somewhat less frequently (Fig. 4). In particular, the selection frequency was high near the center of the northern and southern parts, where the elevation is high. Among the sites selected for the purpose of conserving snow meadow, fellfield, wilderness, and tourism resources, a limited number of sites were selected with high frequency, whereas many sites were selected with low frequency for the conservation of shrubs.

**Fig. 4.**
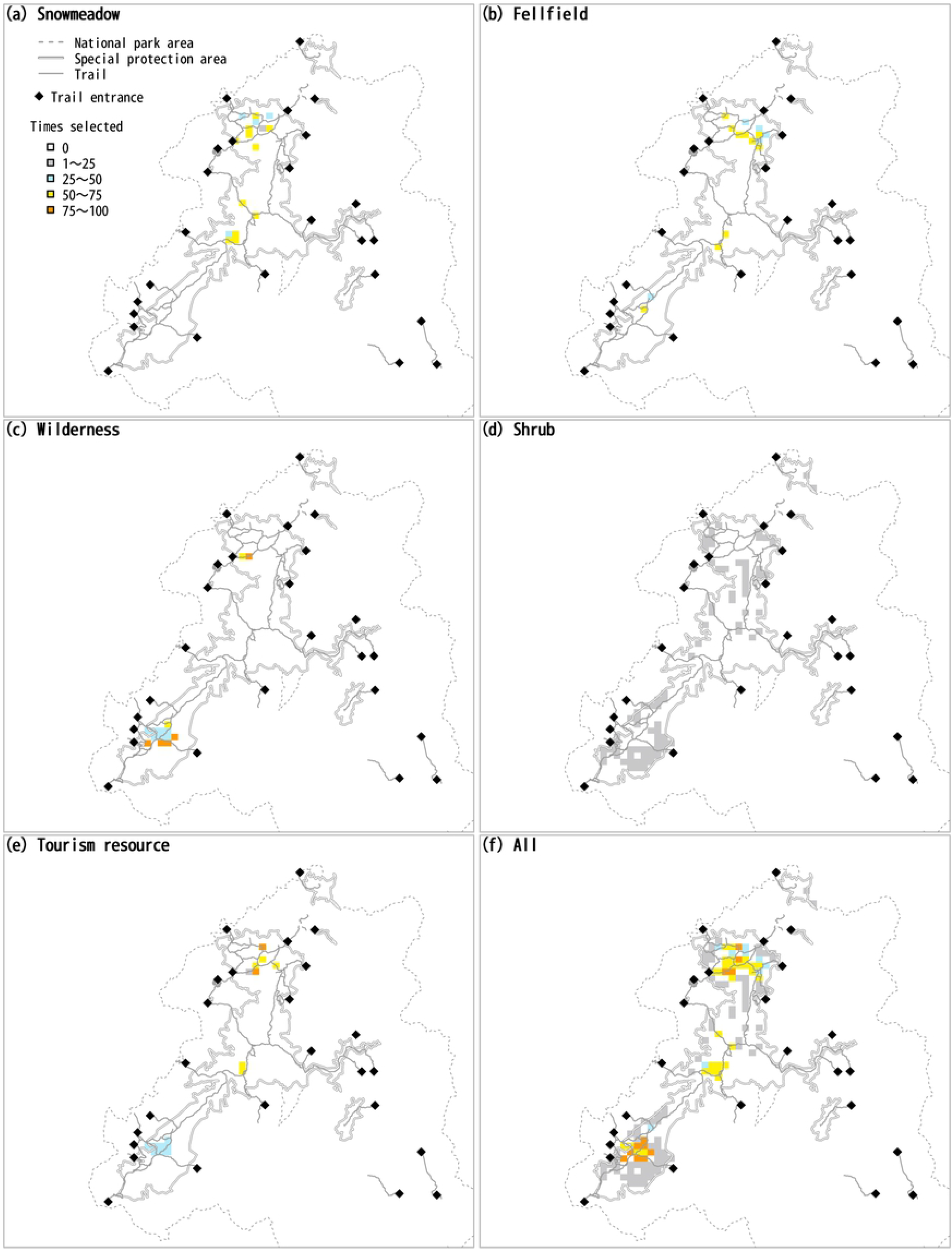
Frequency of selection of each site as areas for conservation of each type of vegetation, for tourism, and for all types of vegetation together.

This pattern of selection likely resulted from the fact that there were many cells where shrubs grew, but the distribution of snow meadow, fellfield, and wilderness was more limited (Fig. 3 and Table 2). Although the target vegetation types overlapped between tourism use and biodiversity conservation, SecSel selected separate sites for these objectives in accordance with the settings when there were conflicts between them. Sites where the area of each type of vegetation was large tended to be selected for conservation, and sites where the total area of the three types of vegetation was large tended to be selected for tourism in accord with the definition of the values of each site for each feature.

**Table 2.** Number of cells in each rank for each type of vegetation and ecosystem service.

| Rank | Snow<br>meadow | Fellfield | Wilderness | Shrub | Tourism<br>resource |
| --- | --- | --- | --- | --- | --- |
| 0 | 201 | 239 | 256 | 44 | 213 |
| 1 | 83 | 36 | 35 | 97 | 53 |
| 2 | 17 | 22 | 13 | 92 | 35 |
| 3 | 17 | 21 | 14 | 85 | 17 |

When consideration was given to cost (cost-on in SecSel), the cost function worked as expected, and the selected sites concentrated close to the trail entrances (Fig. 5 (b)). When the boundary length was taken into consideration (boundary-on in SecSel), the frequency of selection of sites in the central part of the Taisetsu Mountains decreased (Fig. 5 (c)) relative to the frequency with the cost-off and boundary-off settings (Fig. 5 (a)). The highest-quality sites for snow fields and fellfields are in the northern part, and the highest-quality wilderness sites are in the southern part. The northern and southern sites were thus selected in the early steps of the prioritization, and the sites subsequently selected were chosen because of their spatial cohesion around the previously selected sites. The resultant length of the boundary of the selected conservation area averaged 66.8 after 100 runs during which consideration was given to a boundary length penalty. If no consideration was given to the boundary penalty, the lengths of the boundary averaged 110.7 and 97.7 for the cost-off (Fig. 5 (a)) and cost-on (Fig. 5 (b)) settings, respectively. The boundary length of the selected sites was thus substantially decreased when consideration was given to the boundary-length penalty, as expected.

**Fig. 5.**
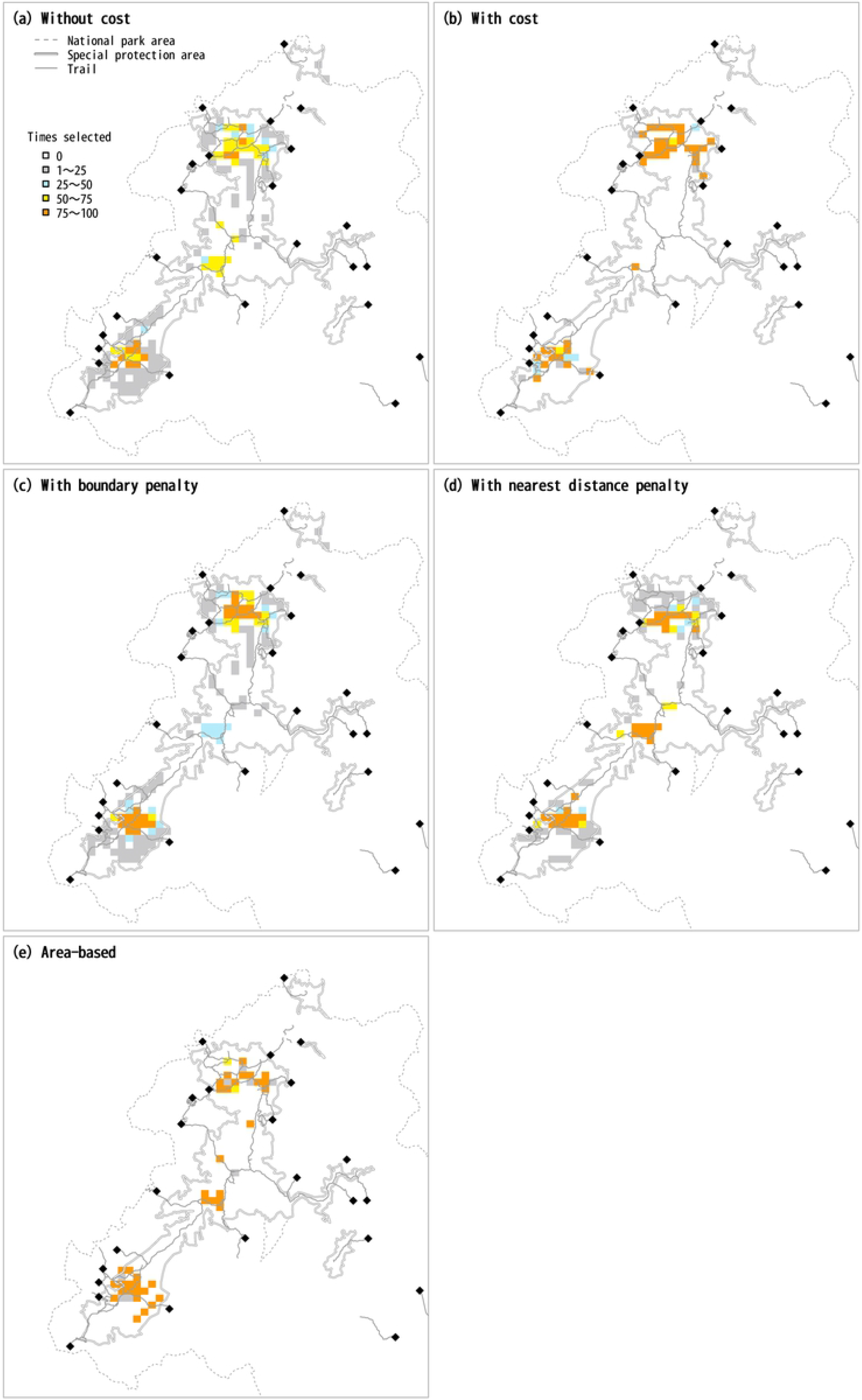
Comparison of site selection results with different SecSel parameter settings.

When consideration was given to the nearest-distance penalty (distance-on), there was an increase in the frequency with which sites in the central part of the Taisetsu Mountains were selected (Fig. 5 (d)), in contrast to the results with the boundary-on setting. Selected sites must adjoin each other to reduce the boundary length, and thus the boundary penalty hindered selection of sites in the central area, which is separate from the northern and southern parts, where there are larger numbers of high-quality sites. The nearest-distance penalty required that a newly selected site be close to already-selected sites, but it was not necessary that the site be adjoint. Furthermore, even larger numbers of sites were selected in the central part to avoid isolation of them and thereby decrease the nearest distance to other sites. The resultant distances to the nearest site averaged 1.99 and 12.90 with and without consideration of the nearest-distance penalty, respectively.

Examination of the site-selection pattern for each run revealed that a small number of sites in the central part were consistently selected on all the example runs when the SecSel settings were cost-off, boundary-off, and distance-off (the first column on the left in S3 Figs.). In contrast, the number of sites selected in the central part varied greatly between runs when the settings were boundary-on or distance-on (the third and fourth columns from the left in S3 Figs.). This difference was caused by the case-dependency of the boundary length and nearest-distance penalties. If a site was selected by chance in the central part in an early step of the site-selection process, sites close to the site were selected to avoid isolation of the site and thereby decrease the boundary length or the nearest-distance to the site. If no site was selected by chance in the central part in the earlier steps, almost no site was selected in the later steps either.

If we used continuous values for site quality, a very limited number of sites was repeatedly selected because SecSel selected sites with the highest quality in order, and there was almost no flexibility when we directly used the area of each type of vegetation. For example, exactly eight sites (the same number as the conservation target) were selected for shrubs. The small flexibility for other types of vegetation was due to the flexibility in the process of resolving conflict between tourism and conservation. When there are conflicts with other conservation objectives, the priority of objectives may change at each iteration. The use of ranks as a measure of site quality increases the flexibility of prioritization by ignoring small differences in areas.

## Discussion

### Assessment of SecSel

In a review of conservation planning tools, Sarkar et al. (2006) [3] have proposed six criteria for the assessment of the tools. Here, we use those criteria to evaluate SecSel.

#### Spatial economy: planning tools designed to select sites should either minimize the cost of sites or maximize the representation of features within cost constraints

SecSel tries to find a minimum set of sites that meet the given multiple targets. If the cost of including each site is given, suppression of the total cost is considered. The algorithm used by SecSel is a greedy algorithm, which is heuristic and does not necessarily find the global minimum solution. However, the difference between the solutions of the greedy algorithm and the global minimum was not large in most cases [15]. Because each site is added in a step-by-step fashion to find the local optimum, the tentative set of selected sites, which expands during the process of site selection, partly fulfills the given target. The process helps determination of how large the selected area should be.

#### Computational efficiency: planning tools should resolve data sets rapidly, particularly if multiple scenarios must be evaluated and stakeholders are involved in real-time negotiations

The greedy algorithm allows SecSel to find a solution rapidly. For example, selection from 3000 sites to satisfy the top-3 of 150 features is repeated 100 within 10 seconds on a desktop PC. In the case of the Taisetsu National Park, it took no more than a few seconds for 100 iterations. The short time required for SecSel to find a solution allows SecSel to be used on-site for discussions in real time.

#### Flexibility: planning tools should allow the incorporation of a wide variety of criteria

The data used to evaluate features to be conserved/used need to be on an ordinal scale, but not necessarily on an interval or ratio scale. SecSel can handle semiquantitative data such as expert judgements. The resolution for the ordinal scale can be set arbitrarily. Furthermore, cost data do not need to be on an interval or ratio scale. The small constraints imposed by the data requirements provide high flexibility that is a distinct advantage of SecSel. In the example of Taisetsu National Park, we showed that SecSel could easily treat the course-time from the nearest trail entrance. That metric has qualitatively different units than the vegetation area metric.

#### Transparency: it should be clear why each site is selected

The logic for site selection in SecSel is straightforward. In every step, it is clear why the newly added site is selected: it makes the largest contribution to the achievement of targets. It is clear which of the features within the site contribute to the achievement of targets. This transparency is an advantage of the heuristic step-by-step approach. Furthermore, the transparency is also apparent in the process of resolving conflicts of features within a site. The logic of the resolution of conflicts is simple: local units that are more dispensable to the achievement of the feature are given lower priority. It is apparent from Fig. 4 that the output of SecSel specifies for what feature(s) a site was selected in the conservation area network, and users can identify the management needed for the site.

#### Genericity: planning tools should solve a variety of problems encountered in practice, using data on any set of biodiversity surrogates, from any type of ecosystem and geographical location

SecSel is not designed to solve specific problems. Any type of feature can be handled, as long as the conservation value of the local units of a feature is evaluated on at least an ordinal scale. SecSel assumes that the focal area consists of multiple ‘sites’. The sites are not necessarily of the same size, nor are they arranged in grids. A set of isolated ‘sites’ such as fragments of ecosystems is acceptable.

#### Modularity: the whole system can be a module, and part of the system can be substituted

The conflict resolution part, cost calculation part, and compactness evaluation part are implemented as separate class objects. These parts can be substituted by user-defined python objects. All input and output data are in the form of a simple text file. Data can be passed to and from other systems easily.

#### Stakeholder involvement

Conservation planning tools are not expected to provide a final solution to be handed to practitioners for implementation. Rather, the tools are expected to be used repeatedly because they reflect stakeholders’ interests, values, and opinions. In the process of using conservation-planning tools, stakeholders can be involved in three stages; *a priori* involvement before the use of the tool, interactive involvement during the use of the tool, and *a posteriori* involvement after the use of the tool [16]. With SecSel, stakeholders can be involved in preparation of data (*a priori* involvement), the repetition of calculations while settings are manipulated (interactive involvement), and interpretation of the results of calculations (*a posteriori* involvement). In the following sections, we describe in more detail how stakeholders can be involved in each stage.

#### A priori involvement

Selection of features to be conserved/used is the first step of *a priori* stakeholder involvement. For conservation of biodiversity features, options are numerus. The feature may be ecosystem types or some taxa. For the conservation of taxa, a decision should be made about which taxonomic group and taxonomic level is to be considered. Furthermore, the feature may be the target of conservation by itself, or a surrogate of something else that is the real conservation target. Stakeholders should be involved in selecting features to be handled with conservation tools.

The criteria for evaluating the features is another point to be considered before the collection of data. Acquiring detailed and/or quantitative data involves cost and effort. The trade-off of the cost and the quality of data should be considered, and an appropriate balance should be found among stakeholders.

Stakeholders should reach an agreement about the appropriate conservation/utilization target for individual features. However, the target may be manipulated interactively, as described later in this section.

If some of the sites of the focal area are designated as protected areas beforehand, SecSel starts site selection with those sites and then adds more. It is up to stakeholders to decide with which sites to start. They may be currently protected areas, or areas of special interest for reasons such as historical, social, or scientific value. Stakeholders are expected to provide a wide range of viewpoints.

#### Interactive involvement

The high computational efficiency of SecSel is due to the heuristic algorithm that supports interactive stakeholder involvement. Stakeholders can try various settings and compare results. Settings to be changed include targets for individual features, conflicts to be considered, cost considerations, pre-reservation of sites, and so on. If the given target is too demanding, a compromise between costs and targets may be explored by manipulating targets. In this process, the consequences of lowering targets should be carefully considered, especially if the lowered target is suspected to be insufficient for sustainable conservation of biodiversity features.

Resolution of the evaluation of local units can also be manipulated to control flexibility and get acceptable solutions. Higher-resolution data with more rank levels lead to a narrower range of variation of sets of sites to achieve targets. This is both an advantage and disadvantage. With higher resolution, full advantage can be taken of detailed data. However, a narrower range of variation of the selection leads to smaller flexibility. Reduction of resolution leads to more ties in the calculation process and can lead to larger variation of the results. An appropriate level of resolution may be found through repeated trials.

Whether or not to incorporate costs and boundary length can be tested in repeated trials with SecSel. These factors may be considered or ignored. The results with or without their consideration will be compared during the interactive involvement process.

### A posteriori involvement

A posteriori stakeholder involvement is the process that occurs after the tool proposes a protected area. Because of the stochasticity in the selection process, SecSel provides multiple sets of sites as land use alternatives. Stakeholders can compare them to find an acceptable solution. Stakeholders can assess the validity of the design from various points of views, which are not explicitly incorporated into the site selection by prioritization tools. The processes of selection by SecSel are straightforward. There is a clear reason for the addition of each site: its contribution to the achievement of targets. This clarity is an advantage in the *a posteriori* stakeholder involvement.

The high calculation efficiency of SecSel allows repeated trials of site selection, so that stakeholders can easily go back to the interactive involvement phase if needed. Interactive and *a posteriori* involvement can be fused.

Sharing and sparing are alternative approaches to the compromise of conservation and land utilization[17, 18]. Sharing in this context means sympatric conservation and utilization. Sparing is the allocation of land area to conservation and utilization separately. In SecSel, sparing can be explicitly considered. Conflicts among local units about features within sites are assessed before site selection, and the conflict is resolved by discarding the local units with lower priority. Since secsel proposes a different resolution for conflicts for each run, stakeholders can choose an acceptable resolution at the stage of a posteriori involvement.

Some prioritization tools resolve conflicts among features in different ways. In Zonation, sites suitable for purposes other than conservation are given negative values so that such sites will likely be omitted from protected areas. Marxan with zones [6], a derivative of Marxan, allocates sites into zones before site selection. The allocation is determined by users of the tool. Different targets are set for different zones. In a zone for conservation of wild animals, for example, targets are the total area of the habitats of focal animals or their population size. If a zone is set for agriculture, the target will be the total area of land suitable for cultivation.

The appropriate balance between features that conserve biodiversity and the benefit obtained from land use depends critically on stakeholders’ interests and values. This dependence is also critical to the resolution of conflicts between features of biodiversity. SecSel does not provide a decision about which feature has higher priority globally. It compares the relative priorities of local units for the achievement of targets of individual features. There is no need to compare different features with a common currency. It is advantageous for stakeholders to reach agreement among themselves.

## Acknowledgements

This research was conducted as a part of the Harmonization with Nature Research Program in the National Institute for Environmental Studies, Japan.

## Supporting Information Captions

**S1 Figs. Site-selection pattern for each run for difference parameter settings for SecSel.**

